# Proteomic analysis of RNA-dependent chromatin association of nuclear proteins

**DOI:** 10.1101/391755

**Authors:** Kyoko Hiragami-Hamada, Naoki Tani, Jun-ichi Nakayama

**Affiliations:** Division of Chromatin Regulation, National Institute for Basic Biology, Okazaki 444-8585, Japan; Department of Basic Biology, School of Life Science, The Graduate University for Advanced Studies (SOKENDAI), Okazaki 444-8585, Japan; Liaison Laboratory Research Promotion Center, Institute of Molecular Embryology and Genetics, Kumamoto University, Kumamoto 860-0811, Japan

**Author notes:** Corresponding authors: Kyoko Hiragami-Hamada (Ph.D.), Assistant Professor, Division of Chromatin Regulation, National Institute for Basic Biology, 38 Nishigonaka, Myodaiji, Okazaki 444-8585, Japan e-mail., Jun-ichi Nakayama (Ph.D.), Professor, Division of Chromatin Regulation, National Institute for Basic Biology, 38 Nishigonaka, Myodaiji, Okazaki 444-8585, Japan.

**Keywords:** RNA-binding proteins, RNA, chromatin, transcription

## Abstract

Various coding and non-coding transcripts are known to associate with chromatin and now there is accumulating evidence that interaction between RNA-binding proteins (RBPs) and RNA molecules regulate not only co-transcriptional mRNA processing, but also other biological processes within the nucleus. Although over a thousand of RBPs have been identified by several mass spectrometry-based methods, it is still unclear which of these RBPs actually associate with chromatin, especially through interaction with RNAs. In addition, biological outcomes of such RBP-RNA-chromatin interactions are yet to be elucidated.

Here we describe a simple proteomics-based method for systematic screening of RBPs that are anchored to chromatin and/or insoluble nuclear substructures by RNA molecules. We used RNase A to release such RBPs from chromatin fraction and analyzed ‘RNase A-solubilized’ proteins by mass spectrometry. Using this method, we were able to identify 156 RNase A-solubilized proteins of which 144 were known RBPs/RBP candidates. Interestingly, several key players of the non-homologous end-joining (NHEJ) pathway were enriched in RNase A-solubilized fraction and the RNA-mediated chromatin association of these factors appeared to be dependent on transcriptional elongation. Furthermore, some enzymes involved in metabolic pathways were also released from chromatin and/or an insoluble nuclear structure by RNase A treatment. In summary, our methodology is highly versatile and is potentially a useful tool to unravel new biological functions for RBP-RNA-chromatin interactions.

## Introduction

Since the discovery of a large collection of mammalian non-coding RNAs in mid-2000, intense research has been carried out around the world to unravel the physical and functional properties of these non-coding transcripts^1,2^. In recent years, it has become apparent that many long non-coding RNAs (lncRNAs) are localized in the nucleus and a pool of chromatin-associated RNAs (caRNAs) have been identified^3,4^. Chromatin act as a platform for many biological processes. The most active event taking place on chromatin is transcription and it was found that a large proportion of caRNAs are products of ongoing transcription^4,5^. These transcripts can recruit factors involved in co-transcriptional processing like splicing of pre-mRNA. More intriguingly, transcription of coding and non-coding transcripts from a specific locus or chromosomal region can, in some cases, recruit chromatin modifiers and/or chromatin binding proteins to mediate epigenetic regulation of gene activity^26,7^. Furthermore, a chromatin-associated lncRNA has been shown to interact with a nuclear matrix protein to induce clustering of individual gene loci *in trans*^8^. However, since information on a functional interaction between a particular caRNA and a certain RNA binding protein (RBP) are rather scarce and fragmentary, it is still difficult to generalize and categorize different types of caRNA – RBP interactions and link them to a repertoire of chromatin-based biological processes.

In parallel to the ‘RNA-centric’ studies, development of crosslinking and immunoprecipitation (CLIP)-mass spectrometry (MS) and similar methodologies greatly accelerated identification of RNA-binding proteins (RPBs)^9-13^. At the present time, over 1,000 proteins have been identified as RBP candidates in mouse and human cultured cells^14^. Conrad *et al.* addressed nuclear localization or enrichment of these RBPs and identified sets of nuclear RBPs and chromatin-associated RBPs (caRBPs)^9, 12^. However, the use of oligo-dT pulldown in the study is expected to have introduced a bias towards identification of RBPs interacting with polyA+ RNAs and may have excluded RBPs associating preferentially to polyA- RNAs, including many nascent transcripts that are the major source of caRNAs. Therefore, it remains unclear which known or unknown RBPs function on chromatin and/or physically interact with caRNAs. Furthermore, various versions of CLIP-MS are powerful methods but, in general, they are not cost-friendly and/or require complicated analyses. This hinders expansion and exploration of RBP research in various biologically relevant contexts.

In order to establish a simple and reliable method to systematically identify caRBPs, especially whose association with chromatin is RNA-dependent, we adopted and modified the method to isolate caRNAs for proteomic study^4,9^. Using HeLa S3 cells, we performed cell fractionation to enrich chromatin and insoluble nuclear substructures. This was followed by RNase A treatment of the chromatin/insoluble fraction to release proteins that are anchored or bridged to chromatin via RNA, and the RNase A-solubilized proteins were analyzed by mass spectrometry. As a result, we identified 156 proteins that were released from chromatin/insoluble nuclear structures by RNase A treatment. The majority of the identified caRBP candidates were already known RBPs or RBP candidates. This method is highly applicable to different types of cells or organisms, thus potentially enables us to examine and explore caRNA-caRPB interactions in different biological contexts.

## Materials and Methods

### Cell culture

HeLa S3 cells were grown in high-glucose DMEM (Nacalai tesque), 10% FBS (Gibco), 1x Penicillin-Streptomycin (Nacalai tesque). To inhibit transcriptional elongation, the cells were treated with 100 μM 5,6-dichloro-1-b-D-ribofuranosylbenzimidazole (DRB) (Sigma Aldrich) for 2 or 4 h.

### Cell Fractionation and RNase A treatment

Cell fractionation was carried out essentially as described by Werner *et al.*^4^. In brief, 0.8 – 1 x 10^7^ HeLa S3 cells were grown in a 10 cm tissue-culture dish and washed twice with ice-cold Dulbecco’s phosphate-buffered saline without Mg^2^+ and Ca^2^+ (D-PBS). The cells were scraped off from the dish and centrifuged at 500 x g for 5 min at 4°C. The supernatant was removed and the cell pellet was resuspended in 2.5x bed volume of Buffer A (10 mM Tris-HCl [pH7.5], 10 mM KCl, 10% glycerol, 340 mM Sucrose, 4 mM MgCl_2_, 1 mM dithiothreitol [DTT], 1x protease inhibitor cocktail (cOmplete, EDTA-free, Roche). Then, an equal amount of Buffer A containing 0.2% Triton X-100 was added. The mixture was incubated for 15 mins on ice and centrifuged at 1,200 x g for 5 min at 4 °C. The resulting nuclear pellet was washed once with 1 ml of Buffer A and then, resuspended in 250 μl of NRB (20 mM Tris-HCl [pH7.5], 50% glycerol, 75 mM NaCl, 1 mM DTT, 1x cOmplete, EDTA-free [Roche]) and centrifuged at 500 x g for 5 min at 4 °C. The nuclear pellet was resuspended in 250 μl of NRB. The equal volume of Buffer NUN (20 mM Tris-HCl [pH7.5], 300 mM NaCl, 1 M Urea, 1% NP-40, 10 mM MgCl2, 1 mM DTT) was subsequently added, incubated for 5 min on ice and centrifuged at 1,200 x g for 5 min at 4 °C. The supernatant was recovered as ‘nucleoplasmic fraction’. The insoluble pellet was washed twice with 1 ml of Buffer A as described above. The washed pellet was resuspended in 250 μl of Buffer A and 100 μl-aliquots were transferred into two 1.5 ml tubes. One aliquot was treated with 5 μg RNase A (10 mg/ml, DNase-free, Nacalai tesque) for 30 min at room temperature. Another aliquot was kept as RNase A-untreated control. After the incubation, the samples were centrifuged at 1,200 x g for 5 min at 4 °C, and the supernatant was collected as ‘RNase A-soluble’ fraction. The pellet was washed once with 1 ml of Buffer A and resuspended in 0.5 ml of Buffer A. To all fractions, 6x SDS sample buffer was added to the final concentration of 1x. The samples were boiled for 5 min at 95 °C, resolved on a 4 – 20% gradient SDS polyacrylamide gel (Cosmobio) and stained with rapid stain CBB kit (Nacalai tesque, for general checks) or GelCode Blue Safe Protein Stain (Thermo Fisher Scientific, for mass spectrometry).

### Mass spectrometry and data analysis

For the in-gel digestion of proteins, each lane was excised 13-16 gel slices, and then these gel slices were cut approximately 1 mm-sized pieces. Proteins in the gel pieces were reduced with DTT (Thermo Fisher Scientific), alkylated with iodoacetamide (Thermo Fisher Scientific), and digested with trypsin and lysyl endopeptidase (Promega) in a buffer containing 40 mM ammonium bicarbonate, pH 8.0, overnight at 37°C. The resultant peptides obtained from in-gel digestion were analyzed on an Advance UHPLC system (AMR/Michrom Bioscience) coupled to a Q Exactive mass spectrometer (Thermo Fisher Scientific) processing the raw mass spectrum using Xcalibur (Thermo Fisher Scientific). The raw LC-MS/MS data was analyzed against the SwissProt or NCBI non-redundant protein/translated nucleotide database restricted to *Homo sapiens* using Proteome Discoverer version 1.4 (Thermo Fisher Scientific) with the Mascot search engine version 2.5 (Matrix Science). A decoy database comprised of either randomized or reversed sequences in the target database was used for false discovery rate (FDR) estimation, and Percolator algorithm was used to evaluate false positives. Search results were filtered against 1% global FDR for high confidence level. The resulting datasets was further analyzed using Scaffold 4 using following cut-off values: minimum number of peptides = 2, peptide threshold = 95%, protein threshold = 99.9%. RNase A-solubilized samples and untreated control samples from 3 experiments were then divided into ‘sample’ and ‘control’ group, respectively. Quantification was carried out using total spectrum counts for each protein. Any ‘sample’ proteins that showed ≥ 2 fold enrichment over ‘control’ and passed a statistical test (Fisher’s exact test in combination with Benjamini-Hochberg procedure) were regarded as RNase A-solubilized proteins. The set of RNase A-solubilized proteins was subsequently subjected to gene ontology (GO), KEGG pathway and InterPro domain enrichment analyses as well as a protein-protein network analysis using STRING (version 10.5, https://string-db.org/). Whole human genome was used as the statistical background for the enrichment analysis. For the network analysis, the confidence level of 0.90 was applied.

### Western blotting

Samples were resolved on a 5 – 20% gradient SDS polyacrylamide gels (Nacalai tesque). The separated proteins were blotted onto PVDF membrane for 1 x~1.5 h at 100 V. The membranes were blocked with 5% skim milk in TBS-T (50 mM Tris-HCl [pH7.6], 150 mM NaCl, 0.05% Tween 20) for 30 min on a rocking platform and then, incubated with primary antibodies (diluted with 2% skim milk in TBS-T) for 2 h at room temperature. The membranes were washed with TBS-T three times (5 min each round on a rocking platform, at room temperature). The washed membranes were incubated with secondary antibodies (diluted in 5% skim milk in TBS-T) for 1 h at room temperature. The membranes were then washed with TBS-T three times (8 ~ 10 min each round on a rocking platform, at room temperature). Signals from the antibodies were detected using ECL (GE Healthcare) or ECL plus reagent (Thermo scientific pierce) and images were captured with LAS-3000 mini (Fuji film). The antibodies used in this study are listed in supplementary table S1.9.

## Results and Discussion

### Enrichment of proteins bound tightly to chromatin

In order to enrich chromatin and other insoluble nuclear substructures including nuclear envelop and nuclear scaffold/matrices, HeLa S3 cells were fractionated into ‘cytoplasmic’, ‘nucleoplasmic’ and ‘chromatin/insoluble’ fractions using the method describe to enrich caRNAs^4^ (Figure 1A). Cells were first lysed in hypotonic buffer containing a non-ionic detergent, Triton X-100, and then, isolated nuclei were lysed in a buffer containing a near physiological concentration (ca. 180 mM) of monovalent salt and a moderate concentration (0.5 M) of urea. The addition of urea to the buffer is thought to effectively reduce RNAs and proteins weakly or non-specifically bound to chromatin^4,15^. As shown in Figure 1B, CBB-staining patterns of each fraction were quite distinct from each other (Figure 1B, lanes 2-4 ‘cytoplasmic’; lanes 6-8 ‘nucleoplasmic’; lanes 10-12 ‘chromatin’). According to western blot analysis of each fraction, typical controls for cytoplasmic and nuclear fractions were enriched in expected fractions; α-tubulin was enriched in cytoplasmic fraction whereas Lamin B and histone H3 were enriched in chromatin/insoluble fractions (Figure 1C). Other nuclear proteins (TIF1ß, HP1β, hnRNP U, Ku80) showed less clear-cut distribution patterns and some of these factors were more enriched in cytoplasmic and nucleoplasmic fractions than in chromatin fraction (Figure 1C). It is likely that Triton X-100 in the hypotonic buffer permeabilized nuclei and this led to release of loosely chromatin-bound and/or abundant nuclear proteins in the ‘cytoplasmic’ fraction. Further treatment with a salt and urea seems to have enhanced release of nuclear proteins associated with nuclear substructures. In summary, the fractionation method seems to have sufficiently removed loosely associated proteins from nuclei therefore allowed us to enrich proteins and RNAs that are stably associated with chromatin/insoluble nuclear substructures.

**Figure 1.**
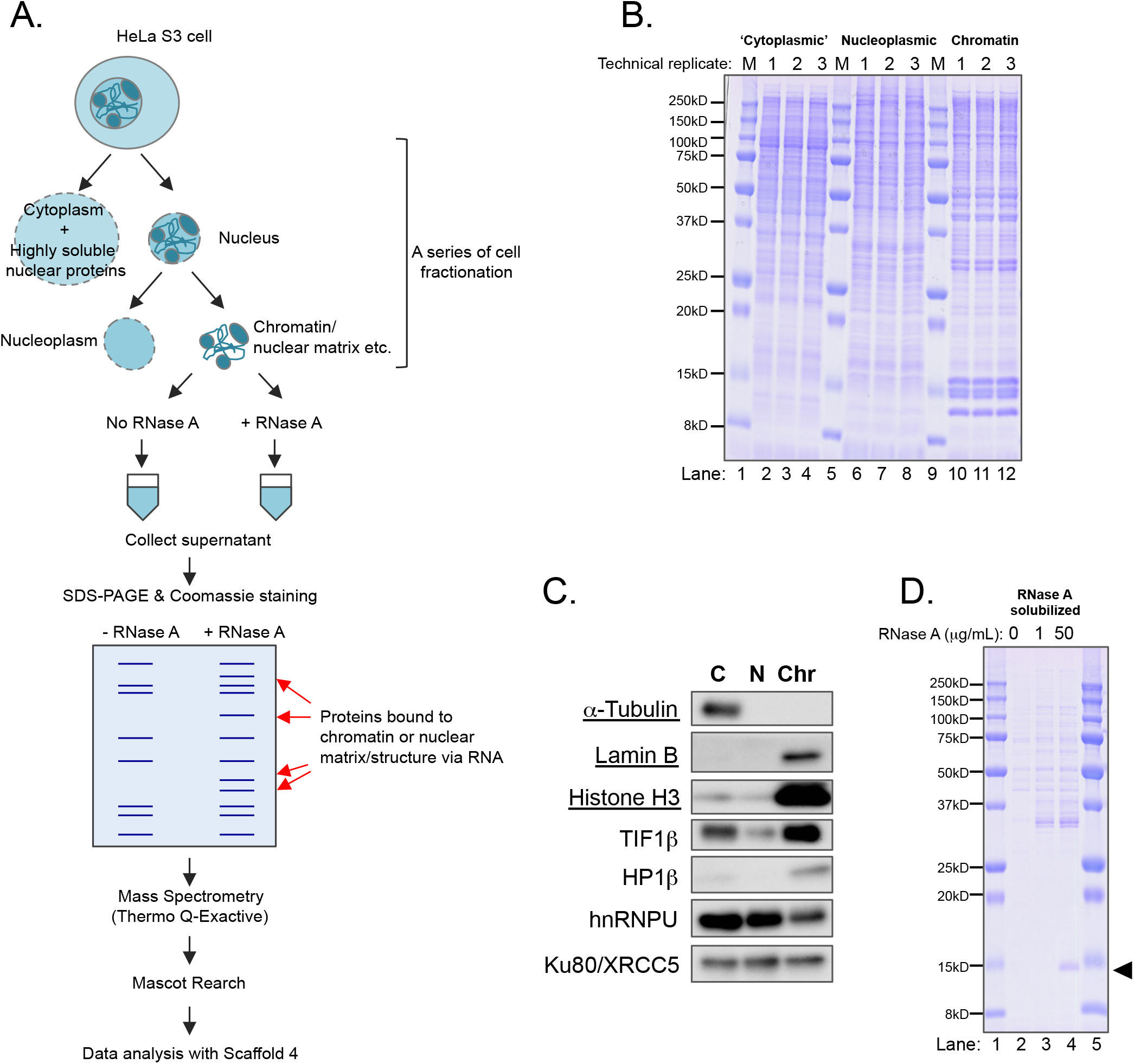
Cell fraction and release of RBPs from chromatin by RNase A treatment. (**A**) A Schematic diagram showing the flow of experimental procedures. (**B**) Proteins from each step of cell fraction and those released by RNase A treatment were analyzed by SDS-PAGE and subsequent CBB staining. ‘M’ stands for a protein size marker. (**C**) Western blot analysis of a selection of proteins in ‘cytoplasmic’(C), ‘nucleoplasmic’ (N) and ‘chromatin/insoluble nuclear substructures’ (Chr). The controls typically used to validate the quality of fractionation are underlined. (**D**) Proteins released from ‘chromatin’ fraction by RNase A treatment. Proteins in supernatants after treatment with various concentrations of RNase A were resolved by SDS-PAGE and visualized by CBB staining. ‘M’ stands for a protein size marker. Black arrow indicates the position of RNase A (visible on lane 4).

### RNase A treatment releases a set of proteins from chromatin fraction

The isolated ‘chromatin’ fraction was then treated with RNase A to solubilize proteins that are anchored to chromatin through interaction with RNA or via association with such RBPs. Under the low salt condition used in this study, RNase A is less specific to single-stranded (ss) RNAs and is expected to digest also double-stranded (ds) RNA molecules to some extent. SDS-PAGE analysis and subsequent CBB staining revealed that a distinct set of proteins were released from the chromatin/insoluble fraction by RNase A treatment (Figure 1D, lane 3, 4), which is evident when compared with untreated control (Figure 1D, lane 2). It seemed that the band intensity and patterns of RNase A-solubilized proteins did not change drastically when treated with 1 μg/ml or 50 μg/ml RNase A (Figure 1D, lane 3, 4). This unresponsiveness to the increase in RNase A dose may indicate that chromatin-associated proteins which can be released by RNase A treatment are rather limited, at least for those appear in the major bands.

### Mass spectrometry analysis of RNase A-solubilized proteins

To identify proteins released from chromatin or insoluble nuclear substructures by RNase A treatment, proteins in the supernatants from RNase A-treated and −untreated samples were analyzed by LC-MS/MS (Figure 1A for scheme, Figure 2A). After several filtering processes, proteins which showed ≥ 2 fold enrichment over untreated controls and passed the statistical test (Fisher’s exact test (p < 0.05) with Benjamini-Hochberg correction; corrected p-value < 0.016161) were regarded as RNase A-solubilized proteins or caRBP candidates and subjected to further analysis. As the result, 156 proteins were identified as caRBP candidates (Supplementary Tables S1.1-1.4). Among 156 proteins, 155 proteins were found in at least 2 out 3 replicates and 136 proteins in all replicates, confirming good reproducibility of identified proteins from 3 independent experiments (Figure 2B). According to GO Molecular Function of the caRBP candidates, the majority of these candidates were annotated RNA- or nucleic acid-binding proteins (Table 1). Also, InterPro Protein Domain analysis of the caRBP candidates indicated that about a third of the proteins contained an RNA-binding motif (Supplementary Tables S2). In agreement with these findings, comparison of the identified proteins against published lists of RBPs/RBP candidates^10,11,13^ also revealed that approximately 92% of them (144/156) were already identified RBPs/RBP candidates (Supplemental Table S1.3). These findings assured that RBPs are preferentially released from chromatin/insoluble nuclear structures and we hereafter refer to the identified proteins simply as caRBPs.

**Figure 2.**
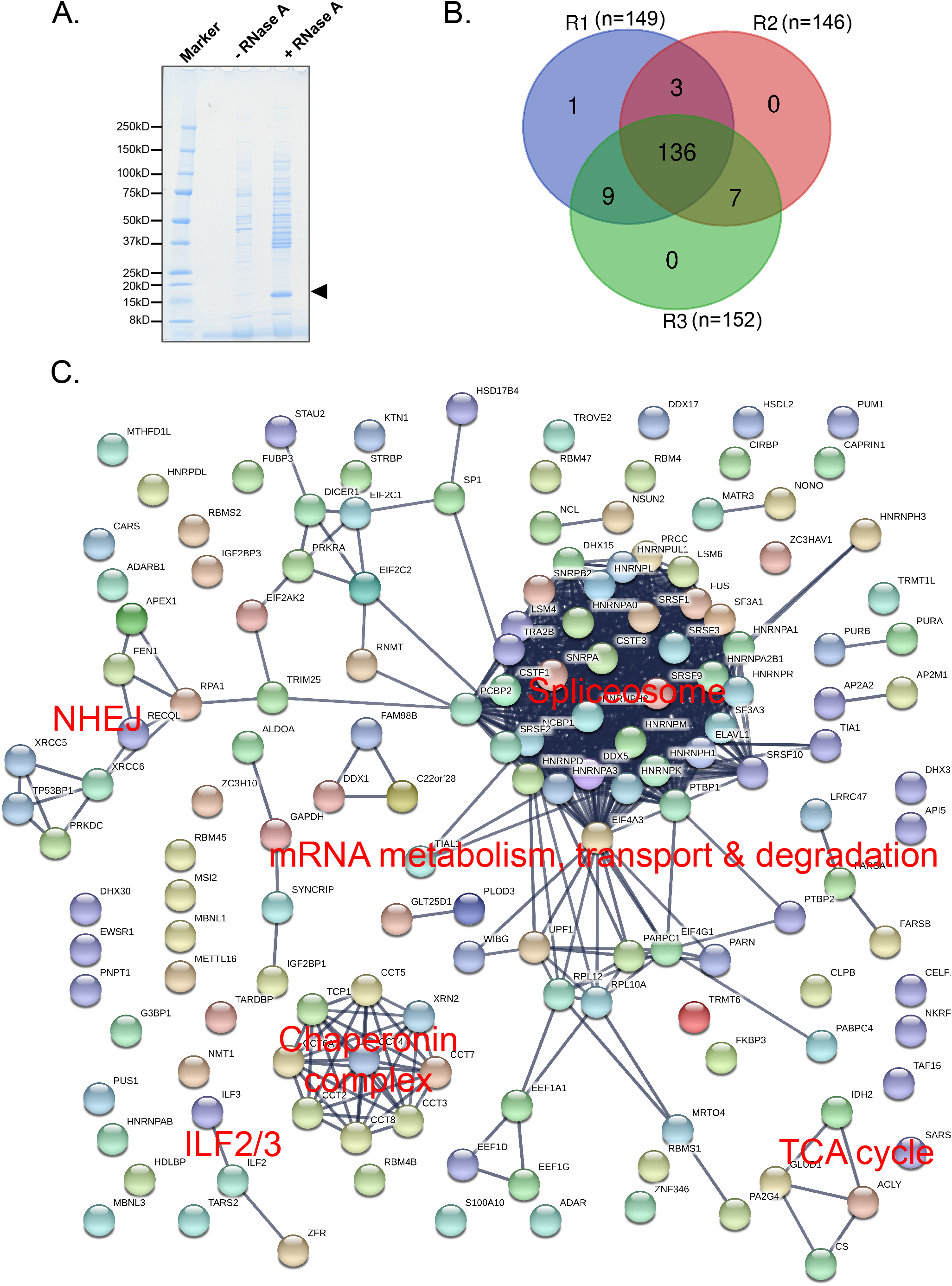
Analysis of RNase A-solubilized caRBPs by mass spectormety. (**A**) An example of RNase A-untreated and −treated samples analyzed by mass spectrometry. Proteins in supernatants after RNase A treatment were resolved by SDS-PAGE and visualized by staining with GelCode Blue dye. Black arrow indicates the position of RNase A. (**B**) A Venn diagram showing overlaps of caRBP candidates from 3 independent experiments. (**C**) STRING network analysis of ‘high confidence’ RNase A-solubilized caRBPs.

**Table 1:**
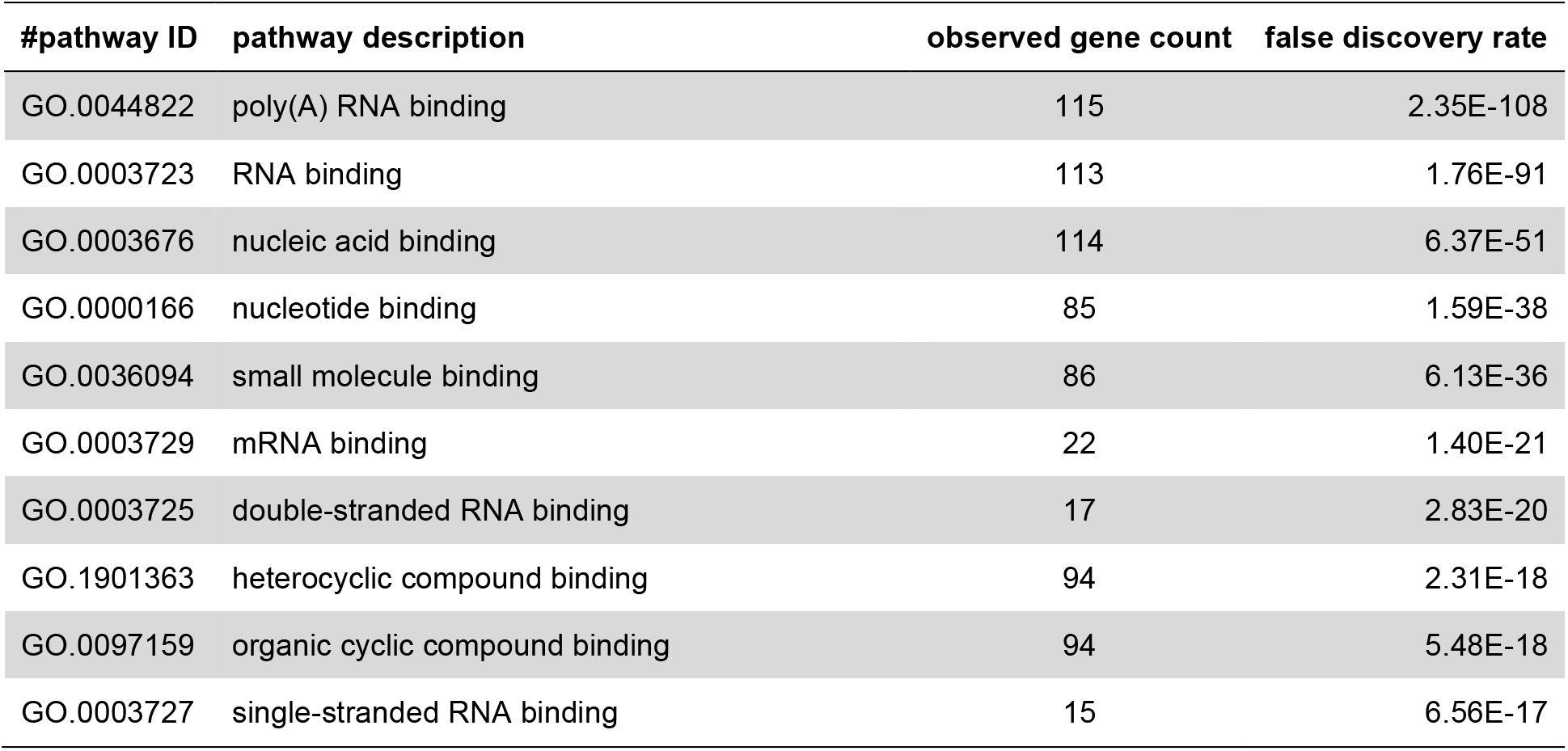
Top 10 GO Molecular Functions of RNase A-solubilized caRBPs. A full, detailed list is provided in Supplementary Tables S2.

| #pathway ID | pathway description | observed gene count | false discovery rate |
| --- | --- | --- | --- |
| GO.0044822 | poly(A) RNA binding | 115 | 2.35E-108 |
| GO.0003723 | RNA binding | 113 | 1.76E-91 |
| GO.0003676 | nucleic acid binding | 114 | 6.37E-51 |
| GO.0000166 | nucleotide binding | 85 | 1.59E-38 |
| GO.0036094 | small molecule binding | 86 | 6.13E-36 |
| GO.0003729 | mRNA binding | 22 | 1.40E-21 |
| GO.0003725 | double-stranded RNA binding | 17 | 2.83E-20 |
| GO.1901363 | heterocyclic compound binding | 94 | 2.31E-18 |
| GO.0097159 | organic cyclic compound binding | 94 | 5.48E-18 |
| GO.0003727 | single-stranded RNA binding | 15 | 6.56E-17 |

As expected, GO Biological Function and KEGG pathway analysis of the identified caRBPs revealed that a large proportion of these proteins are related to mRNA biology including mRNA processing, surveillance, metabolism, transport and degradation (Figure 2C, Table 2, 3). Besides this expected observation, some unexpected and interesting groups of proteins were identified. In particular, proteins involved in the non-homologous end-joining (NHEJ) DNA repair pathway (DNA-PKcs, Ku80/XRCC5, Ku70/XRCC6, FEN1) and several metabolic enzymes, especially those involved in the tricarboxylic acid (TCA) cycle (ACLY, CS, IDH2) were markedly and consistently enriched in RNase A-solubilized fraction (Figure 2C, Table 3). The presence of metabolic enzymes in chromatin-enriched fraction could be mere contamination by mitochondria and/or cytoplasmic materials. However, recent studies indicate that metabolic enzymes can indeed shuttle between mitochondria and nucleus and those in the nucleus may participate in transcriptional/chromatin regulation^16^. Also, it must be noted that the RNA-binding property of metabolic enzymes has long been known^17,18^. Together with these circumstantial evidences, our finding supports the idea that a set of metabolic enzymes is associated with chromatin and raises the possibility that such interaction involves RNA.

**Table 2.** Top 10 GO Biological Functions of RNase A-solubilized caRBPs. A full, detailed list is provided in Supplementary Tables S2.

| #pathway ID | pathway description | observed gene count | false discovery rate |
| --- | --- | --- | --- |
| GO.0006396 | RNA processing | 64 | 7.39E-50 |
| GO.0016071 | mRNA metabolic process | 54 | 6.14E-42 |
| GO.0006397 | mRNA processing | 47 | 3.88E-40 |
| GO.0008380 | RNA splicing | 42 | 1.37E-36 |
| GO.0000398 | mRNA splicing, via spliceosome | 35 | 5.22E-34 |
| GO.0016070 | RNA metabolic process | 90 | 1.77E-30 |
| GO.0090304 | nucleic acid metabolic process | 93 | 7.82E-29 |
| GO.0010467 | gene expression | 92 | 4.64E-28 |
| GO.0006139 | nucleobase-containing compound metabolic process | 96 | 1.44E-27 |
| GO.0006725 | cellular aromatic compound metabolic process | 97 | 5.16E-27 |

**Table 3.** KEGG Pathways over-represented by RNase A-solubilized caRBPs. A detailed list is provided in Supplementary Tables S2.

| #pathway ID | pathway description | observed gene count | false discovery rate |
| --- | --- | --- | --- |
| 3040 | Spliceosome | 19 | 2.76E-17 |
| 3015 | mRNA surveillance pathway | 10 | 1.44E-07 |
| 3018 | RNA degradation | 8 | 6.16E-06 |
| 3450 | Non-homologous end-joining | 4 | 1.54E-04 |
| 3013 | RNA transport | 8 | 1.16E-03 |
| 970 | Aminoacyl-tRNA biosynthesis | 4 | 1.49E-02 |
| 5168 | Herpes simplex infection | 7 | 1.51E-02 |
| 20 | Citrate cycle (TCA cycle) | 3 | 4.36E-02 |

In addition to the software-assisted analyses, manual extraction of proteins known to be a part of nuclear substructures or a transcription regulator showed enrichment for several transcription activators/regulators including ILF2/3 and PurA/B in RNase A-solubilized fraction (Table 4). Enrichment for a subset of paraspeckle proteins^19^ was also observed. However, the observation needs to be interpreted with caution since many of the identified paraspeckle proteins are hnRNPs or those with additional functions besides paraspeckle formation. In addition, there was no clear enrichment for a particular class of paraspeckle proteins^19^ (subclasses are indicated in Table 4).

**Table 4.** A list of manually extracted caRBPs implicated in nuclear substructures or transcriptional regulation. ‘INF’indicates proteins only found in RNase A-treated samples but not in controls. Subclasses of paraspeckle proteins are shown in brackets.

| Protein Name | p-value | Fold enrichment | Subnuclear structures/Functions |
| --- | --- | --- | --- |
| CIRBP | 0.0016 | INF | Paraspeckle (3B) |
| EWSR1 | 0.0023 | INF | Paraspeckle (3B) |
| RBM4B | < 0.00010 | INF | Paraspeckle (3B) |
| HNRNPA1 | < 0.00010 | 15 | Paraspeckle (2) |
| HNRNPUL1 | < 0.00010 | 11 | Paraspeckle (2) |
| FUS | < 0.00010 | 8.1 | Paraspeckle (1B) |
| NONO | 0.0031 | 7.8 | Paraspeckle (1A) |
| HNRNPR | < 0.00010 | 5.4 | Paraspeckle (2) |
| TARDBP | < 0.00010 | 5.1 | Paraspeckle (2) |
| HNRNPH3 | < 0.00010 | 4.8 | Paraspeckle (1B) |
| TAF15 | 0.00022 | 4.5 | Paraspeckle (3B) |
| HNRNPH1 | < 0.00010 | 3.4 | Paraspeckle (unclassified) |
| NCL | < 0.00010 | 6.8 | Nucleolus |
| ILF2 | < 0.00010 | 17 | Transcriptional regulator |
| ILF3 | < 0.00010 | 16 | Transcriptional regulator |
| PurA | < 0.00010 | INF | Transcriptional regulator |
| PurB | < 0.00010 | INF | Transcriptional regulator |
| NKRF | < 0.00010 | INF | Transcriptional regulator |

While analyzing the datasets, we realized there were a considerable number of proteins that showed ≥ 2 fold enrichment over the control but failed to pass the statistical test. The primary reasons for this appear to be the high background level of a protein in the control and/or low spectrum/peptide counts for a protein. This group of proteins included several know RBPs such as hnRNP U, hnRNP UL2 and SF1 whose functions are splicing-related and/or whose RNase A-dependent release from chromatin have been demonstrated previously^20^. In fact, western blot analysis of hnRNP U in RNase A-solubilized fractions clearly indicated that the protein had been released from chromatin by the nuclease treatment (Figure 3). Thus, to minimize ‘false negative’ proteins, we manually extracted proteins that sufficed the following criteria; 1) exhibit at least ≥ 2-fold enrichment over corresponding control in at least two replicates and 2) represented by at least two exclusively unique peptides. These proteins were defined as ‘low confidence’ caRBPs. The procedure resulted identification of 179 low confidence caRBPs (Supplementary Table S1.5). Unlike the ‘high confidence’ counterpart, many of them were not annotated for RNA-binding (59/179 [33.0%] for low confidence caRBPs vs 132/156 [84.6%] for high confidence caRBPs). Because the annotation is mainly based on the datasets of PolyA+ RNA-binding proteins generated by Castello *et al.* and Baltz *et al.*^10,11^, we also compared our list with the dataset produced by Bao *et al.*^13^ which include RBPs that interact with PolyA-RNAs and nascent transcripts. This increased the overlap with known RBPs to 124/179 (69.3%) for low confident caRBPs and 144/156 (92.3%) for high confidence caRBPs (Supplmentary Table S1.5). GO Molecular Function and Biological Function analysis suggested that the low confidence caRBPs were also enriched for proteins implicated in RNA-binding and/or RNA biology but, with much higher FDRs compared with high confidence caRBPs (Supplementary Tables S2). Apart from additional splicing factors, KEGG pathway and STRING network analysis of low confidence caRBPs showed distinct clusters of proteins from those of high confidence caRBPs (Supplementary Figure S2, Supplementary Tables S2). Interestingly, proteasome components, SUMOylation-related proteins (SUMO1, SUMO2, UBE2I) and chromatin binding/modifying proteins (SETD3, SMARCC1, SMARCA5, RCOR1, RBBPs, histone H1 and HMGB1) were among the low confidence caRBPs and formed clusters (Supplementary Figure S2). RNA-mediated interactions between these proteins and chromatin are likely to be biologically relevant and may worth further investigation. However, since we noticed that a fraction of the low confidence caRBPs seemed to correspond to proteins from contaminating organelles or cellular fractions. Consequently, we decided to focus on high confidence caRBPs (and hnRNP U from the low confidence candidates) for further analyses.

**Figure 3.**
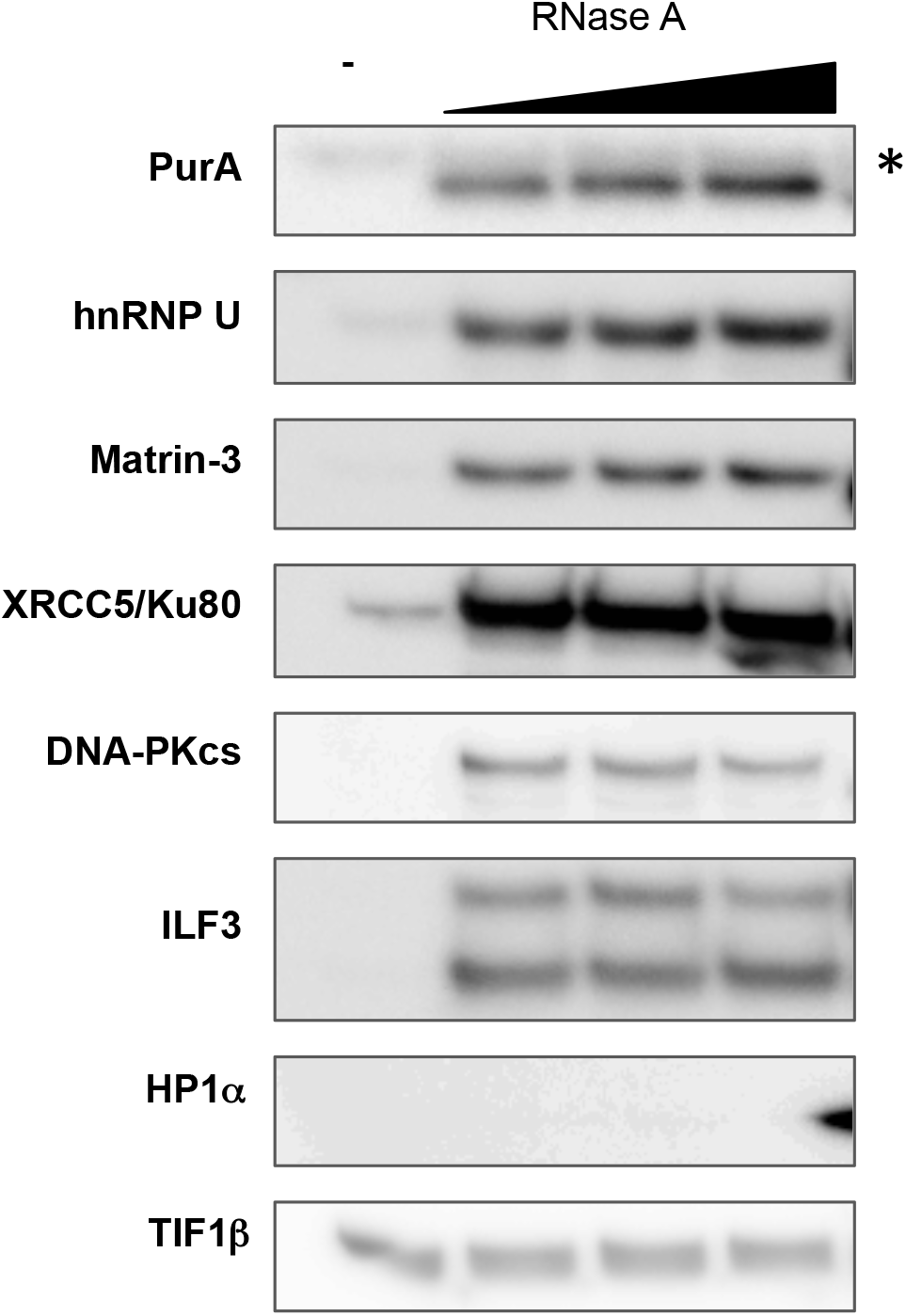
Validation of RNase A-solubilized caRBP candidates by western blotting. A selection of caRBP candidates identified by mass spectrometry were subjected to western blot analysis. RNase A solubilized fraction resulting from treatment with different concentrations of RNase A were resolved by SDS-PAGE, blotted onto PVDF membranes which were subsequently incubated with antibodies against the indicated proteins. Asterisk indicates a non-specific band. HP1α and TIF1β serve as negative controls whose enrichment in RNase A-solubilized fraction was not detected by mass spectrometry.

### Verification of MS results by western blotting

Having obtained the list of RNase A-solubilized caRBPs, a handful of high confidence caRBPs (PurA, ILF3, DNA-PKcs, Ku80/XRCC5, Matrin-3) and hnRNP U were selected and the mass spectrometry results were confirmed by western blot (Figure 3). For all the proteins tested, we could clearly detect increases in signals from the proteins released from chromatin upon RNase A treatment (Figure 3). Interestingly, the levels of PurA, hnRNP U and matrin-3 released from chromatin seemed to be dependent on the RNase A dose to a certain extent; more proteins were released at higher concentrations of RNase A (Figure 3). This increase was more or less concomitant with decrease in chromatin-bound fraction (Supplementary Figure S1). On the other hand, other candidates (ILF3, Ku80, DNA-PKcs) did not show such response to RNase A dose and the majority of these proteins remained on chromatin or insoluble structures even when treated with the highest dose of RNase A (Figure 3 and Supplementary Figure S1). These observations may imply that association of PurA, hnRNP U and matrin-3 with chromatin or insoluble nuclear substructures is mainly RNA-dependent whereas only a fraction of the other proteins interact with chromatin through association with RNA. The majority of these proteins may interact with chromatin or insoluble nuclear substructures by another mode (such as direct interaction with DNA, histones or matrix proteins). Furthermore, consistent with our mass spectrometry results, RNase A-dependent release of HP1α and TIFβ, which are chromatin proteins previously implicated in RNA binding^11,21-23^, was not detected by western blot (Figure 3). In fact, none of classical chromatin binding or modifying factors including polycomb protein ^24-28^ and SUV39H1/2 histone methyltransferases^29-31^ whose RNA-binding activity have been demonstrated *in vitro* and in cultured cells, was identified as caRBPs by our method (Supplementary Tables S1.1-1.5). This could be due to the limited sensitivity of the method. Alternatively, the failure to detect these proteins in RNase A-solubilized fraction could be due to a biological reason. For example, chromatin-binding proteins like polycomb proteins have multiple modules for chromatin interaction, and in the absence or upon removal of RNA, these proteins remain bound to chromatin through interaction with DNA or histone proteins^32-34^. In addition, RNA-dependent recruitment and anchoring of some chromatin factors to chromatin may be prominent only at a certain developmental stage^35^. Nonetheless, our analysis of low confidence caRBPs suggested interaction between several chromatin-related proteins (see the previous subsection) and caRNAs. Moreover, our preliminary analysis of RNase III-solubilized proteins resulted in identification of quite a distinct set of caRBPs, compared with RNase A-solubilized caRBPs (Supplementary Figure S3, Supplementary Tables S1.6-1.8, S2). Curiously, these included several chromatin remodelers (SMARCA5, SMARCC2), histone demethylases (KDM1A, KDM3B) and a chromatin binding protein (AHNAK, TRP). Since RNase III specifically digests dsRNAs, these chromatin factors may preferentially interact with RNA duplexes or RNA molecules with extensive secondary structures.

### Different caRBPs appear to show different sensitivity to transcriptional inhibition

As revealed by recent analysis of global RNA-chromatin interaction, most of caRNAs appear to be nascent transcripts^5^. Thus, we wondered if association between RNA and caRBP is coupled with transcription. In order to address this point, HeLa S3 cells were treated with DRB, a reversible inhibitor of transcriptional elongation, and then subjected to cell fractionation and RNase A treatment (Figure 4A). RNase A-solubilized proteins were subsequently analyzed by western blot. DRB treatment reduced the level of the elongation-associated form of RNA polymerase II (RNAPII Ser2P)^36^ and to some extent overall RNAPII level in the nucleus (Figure 4B), indicating transcriptional elongation is sufficiently inhibited by the reagent. On the other hand, the nuclear protein levels of the caRBPs we tested were not greatly affected by the inhibitor treatment (Figure 4B, Whole nucleus). Having this in mind, the effect of DRB on RNase A-mediated release of the caRBPs seemed to vary from protein to protein. Transcriptional regulators, ILF3 and PurA showed no marked reduction in RNase A-solubilized fraction whereas hnRNP U and matrin-3, which are both nuclear scaffolding factors, showed moderate reduction in RNase A-solubilized fraction upon DRB treatment (Figure 4B). Most strikingly, RNase A-dependent release of the key players of the NHEJ pathway, Ku80 and DNA-PKcs from chromatin was greatly reduced by inhibition of transcriptional elongation (Figure 4B). Although the number of proteins tested is small, these findings indicate that the ‘DRB-sensitivity’ of RBPs may somewhat reflect their functional categories. Transcriptional regulators like ILF3 and PurA might interact with RNA during the initiation stage of transcription and/or associate with RNA in a certain structure/conformation. As a matter of fact, it has been shown that ILF3 (and ILF2) preferentially bind to RNA:DNA hybrids (so-called R-loops) that are enriched around transcriptionally active/permissive promoters on the genome^37^. Regarding the high DRB-sensitivity of the NHEJ factors, the observation agrees with the previous finding that these factors associate with nascent transcripts^13^. Moreover, several recent studies demonstrated that the NHEJ factors are recruited to sites of ongoing transcription^38^ and participate in transcription-coupled DNA repair^39^ or paused RNAPII release for transcriptional elongation^38^. Although we cannot rule out the possibility that the interaction between nascent RNAs and Ku80 or DNA-PKcs is indirect, lack of correlation between the level of RNase A-solubilized RNAPII (general or S2P) and that of RNase A-solubilized Ku80/DNA-PK (Figure 4B) suggests that these NHEJ factors are recruited to and associate with transcriptionally active genomic regions independent of RNAPII, through direct interaction with elongating, nascent transcripts. Taken together, these observations indicate that caRBPs – caRNAs interactions are not uniform and the nature of the interaction might be governed by transcriptional stage, conformation/presentation of caRNAs and functional properties of RBPs on chromatin.

**Figure 4.**
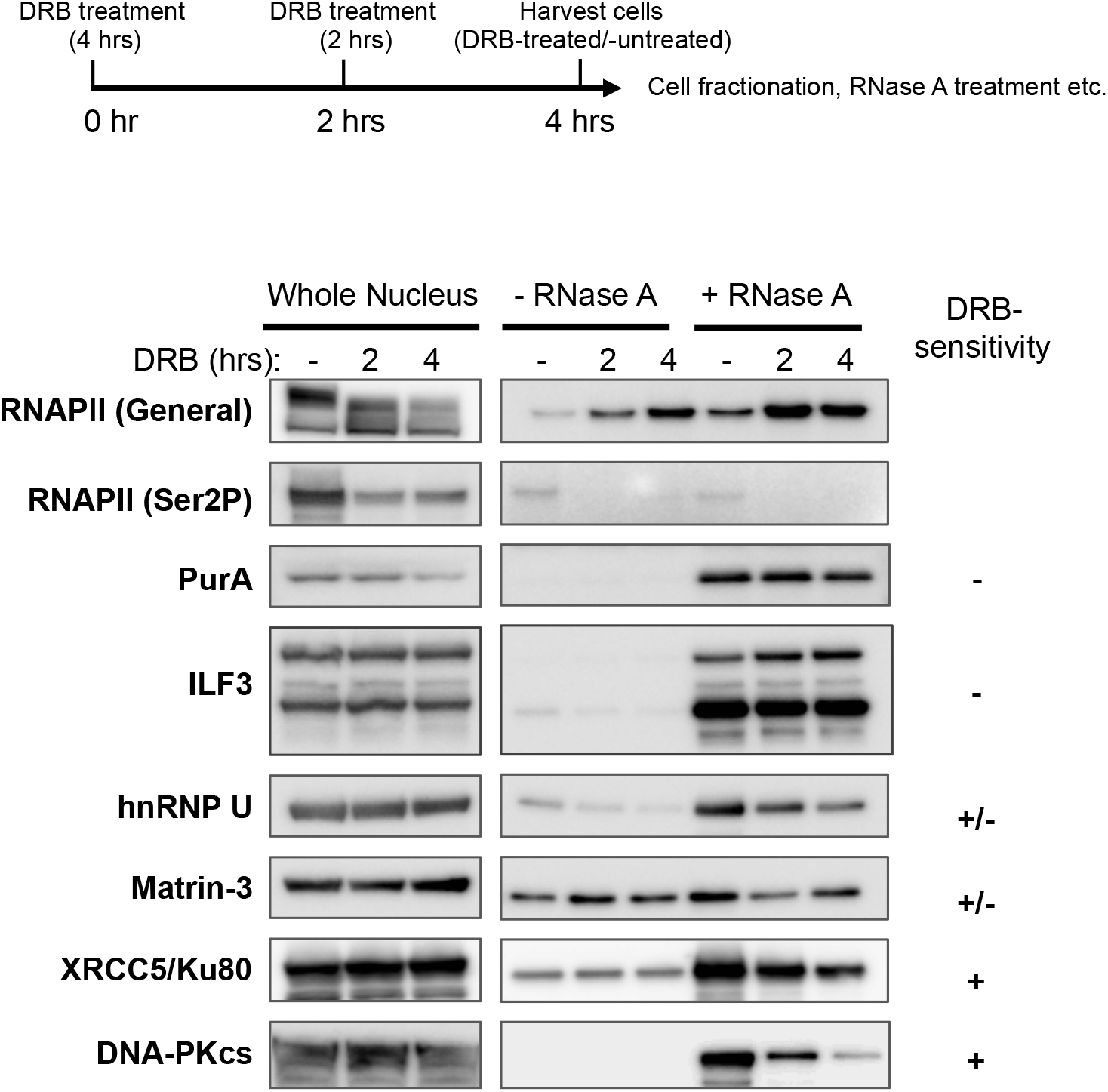
Western blot analysis of RNase A-solubilized caRBP candidates upon inhibition of transcriptional elongation. (**A**) A timeline of DRB treatment. In order to inhibit transcriptional elongation, HeLa S3 cells were treated with 100 μM DRB for 0, 2 or 4 h. After the incubation with DRB, the cells were harvested and subjected to cell fraction and RNase A treatment as in ‘Materials and Methods’ section. (**B**) Western blot analysis of whole nuclear extracts and RNase A solubilized fractions from DRB-untreated and -treated cells. Western blotting was performed with antibodies specific for the proteins indicated.

## Conclusions

In this study we successfully narrowed down caRBPs to approximately 150 among over a thousand of known RBPs using a non-CLIP method. Since our experimental conditions and the cutoff values for analysis were set to identify highly reproducible caRPBs, the caRBPs listed here are likely to be the *‘bona fide*’ caRBPs that participate in the most fundamental biological processes like transcription, DNA repair and mRNA processing, and metabolisms. In order to capture minor caRBPs, further optimization of experimental conditions and/or extra step to enrich such proteins from RNase A-solubilized fraction might be required. In addition, our analyses of RNase A- and RNase III-solubilized caRBPs suggest that use of different RNases may help to release and identify ‘minor’ or additional caRBPs.

Overall, this method is relatively simple, cheap and highly adaptable to any kinds of cells or organisms. By applying this method to different cell types or cells undergoing different stages of a physiological change, it may be possible to systematically identify functionally relevant, cell type-specific or stage-specific caRBPs. Moreover, our results from the DRB experiments suggest that combinatorial use of this methodology with various transcription/splicing inhibitor treatments may enable us to further characterize different types of caRNA-caRPB interactions. Last but not least, it is important to note that integration of proteomics data with genomics/transcriptomics data such as CLIP-seq results of caRBPs is necessary to generate a more comprehensive picture of caRNA-caRPB interactions. In conclusions, we anticipate future applications of this method in different biological as well as experimental contexts will give us deeper insights into caRNA-caRPB interactions.

## Supporting information

**Supplementary Figure S1.** Western blot analysis of chromatin-bound fractions after RNase A treatment.

**Supplementary Figure S2.** STRING network analysis of ‘low confidence’ RNase A-solubilized caRBPs.

**Supplementary Figure S3.** Mass spectrometry analysis of RNase III-solubilized caRBPs. **Supplementary Table S1.** Lists of proteins and spectrum counts generated by mass spectrometry analyses, and a list of antibodies used in this study:

*Supplementary Table Sl.l.* A full list of proteins and total spectrum counts in RNase A-treated and untreated samples.

*Supplementary Table S1.2.* A full list of proteins and unique peptide counts in RNase A-treated and untreated samples.

*Supplementary Table S1.3.* A list of RNase A-solubilized caRBP candidates.

*Supplementary Table S1.4.* A list of RNase A-solubilized caRBP candidates in each experimental replicate.

*Supplementary Table S1.5.* A list of ‘low confidence’ RNase A-solubilized caRBP candidates.

*Supplementary Table S1.6.* A full list of proteins and total spectrum counts in RNase III-treated and untreated samples.

*Supplementary Table Sl. 7.* A full list of proteins and unique peptide counts in RNase III-treated and untreated samples.

*Supplementary Table S1.8.* A list of RNase III-solubilized caRBP candidates.

*Supplementary Table S1.9.* A list of antibodies used in this study.

**Supplementary Table S2:** Summary of GO analyses of RNase A- or RNase III-solubilized caRBP candidates.

*N.B.* ‘A_high’, ‘A_low’ and ‘III’ stand for RNase A-solubilized <u>high</u> confidence caRBPs, RNase A-solubilized <u>low</u> confidence caRBPs and RNase III-solubilized (high confidence) caRBPs, respectively.

## Data availability

Raw data files from mass spectrometry of RNase A-treated and -untreated samples have already been deposited on jPOST database^40^ (https://jpostdb.org; accession numbers: JPST000468 for jPOST and PXD010642 for ProteomeXchange) but are not yet released to general public. For time being, the data files are available from the corresponding authors (K.H-H, J.N) upon reasonable request.

## Author contributions

K.H-H conceived, designed and performed experiments, carried out data analysis with Scaffold 4 and STRING software and wrote the paper. N.T. processed samples for mass spectrometry, performed mass spectrometry and analyzed the data. J.N coordinated the project and wrote the paper.

## Acknowledgements

We would like to thank the Liaison Laboratory Research Promotion Center (Kumamoto University) for technical support. This work was supported by Mochida memorial foundation and Sumitomo foundation (to K.H-H).

